# Structural Dynamics of Real and Modeled *Solanum* Stamens: Implications for Pollen Ejection by Buzzing Bees

**DOI:** 10.1101/2021.10.25.465809

**Authors:** Mark Jankauski, Riggs Ferguson, Avery Russell, Stephen Buchmann

## Abstract

An estimated 10% of flowering plant species conceal their pollen within tube-like anthers that dehisce through small apical pores (poricidal anthers). Bees extract pollen from poricidal anthers through a complex motor routine called floral buzzing, whereby the bee applies vibratory forces to the flower stamen by rapidly contracting its flight muscles. The resulting deformation depend on the stamen’s natural frequencies and vibration mode shapes, yet for most poricidal species these properties have not been sufficiently characterized. We performed experimental modal analysis on *Solanum elaeagnifolium* stamens to quantify their natural frequencies and vibration modes. Based on morphometric and dynamic measurements, we developed a finite element model of the stamen to identify how variable material properties, geometry and bee weight could affect its dynamics. In general, stamen natural frequencies fell outside the reported floral buzzing range, and variations in stamen geometry and material properties were unlikely to bring natural frequencies within this range. However, inclusion of bee mass reduced natural frequencies to within the floral buzzing frequency range and gave rise to an axial-bending vibration mode. We hypothesize that floral buzzing bees exploit the large vibration amplification factor of this mode to increase anther deformation, which may facilitate pollen ejection.

## 1. Introduction

Pollen is an essential resource for plant reproduction as well as the nourishment of approximately 20,000 species of bees globally. Over the past 135-million-years, thousands of animal-pollinated angiosperms have evolved to “dose” the amount of pollen available to an individual floral visitor (Harder & Thomson, 1989; Harder & Wilson, 1994; Percival, 1955). Dosing limits the amount of pollen available during any single floral visit, thus facilitating the dispersal of pollen via multiple pollinators, which can enhance floral reproductive success. One particularly common dosing strategy involves shielding pollen in specialized tube-like anthers that dehisce only via small apical slits, pores, or valves, through which the pollen can be removed (i.e., ‘poricidal anthers’; Buchmann, 1983; De Luca & Vallejo-Marín, 2013). Numerous bee species extract pollen from poricidal anthers by grasping the anther with their mandibles and rapidly contracting their indirect flight muscles, a behavior termed floral buzzing (Cardinal et al., 2018; Harder & Barclay, 1994; Switzer et al., 2019; Vallejo-Marín, 2019). The ensuing vibration of the anthers results in the ejection of pollen grains onto the bee’s body where they can be collected. The successful extraction of pollen relies both on the vibratory forces produced by the bee, as well as the dynamical properties of the poricidal anther and supporting filament. Despite considerable efforts to characterize the vibration of poricidal anthers (e.g., Corbet et al., 1988; Corbet & Huang, 2014; Harder & Barclay, 1994; King & Buchmann, 1995, 1996; M. J. King & L. Lengoc, 1993; Nevard et al., 2021; Nunes et al., 2021, also see Vallejo-Marín, 2019, 2021 and references within), certain biophysical properties such as stamen natural frequencies and vibration mode shapes remain understudied in many poricidal flower species.

The natural frequencies and damping ratios of the stamen (a structure consisting of a pollen-laden anther supported by a narrow filament, Figure 1) may contribute to its deformation during floral buzzing. In general, exciting a structure at one or more of its natural frequencies will elicit a large vibrational response due to resonance, particularly if the damping is low (De Langre, 2019). Stamen resonance plays a critical role in pollen expulsion in some wind-pollinated flowers (Timerman et al., 2014; Timerman & Barrett, 2018, 2019; Urzay et al., 2009). Within poricidal anthers, resonant or near resonant responses would increase kinetic energy transferred from the vibrating locule walls to the pollen particles, which may facilitate pollen expulsion through increased motion (Buchmann & Hurley, 1978). Indeed, amplitude of the vibrating anther has been positively correlated with the amount of pollen expelled in two *Solanum* species (De Luca et al., 2013; Rosi-Denadai et al., 2020). Additionally, the first natural frequency of poricidal anthers from a variety of plant taxa has been measured (e.g., Harder & Barclay, 1994; King & Buchmann, 1996; Nunes et al., 2021, also see Vallejo-Marín, 2019, 2021 and references within). However, continuous structures such as stamens have many natural frequencies (Rao, 2019), and if excited during floral buzzing, these higher-order natural frequencies could play an important role in pollen expulsion. Yet to our knowledge, the natural frequencies beyond the first have not been quantified for poricidal flowers.

**Figure 1.**
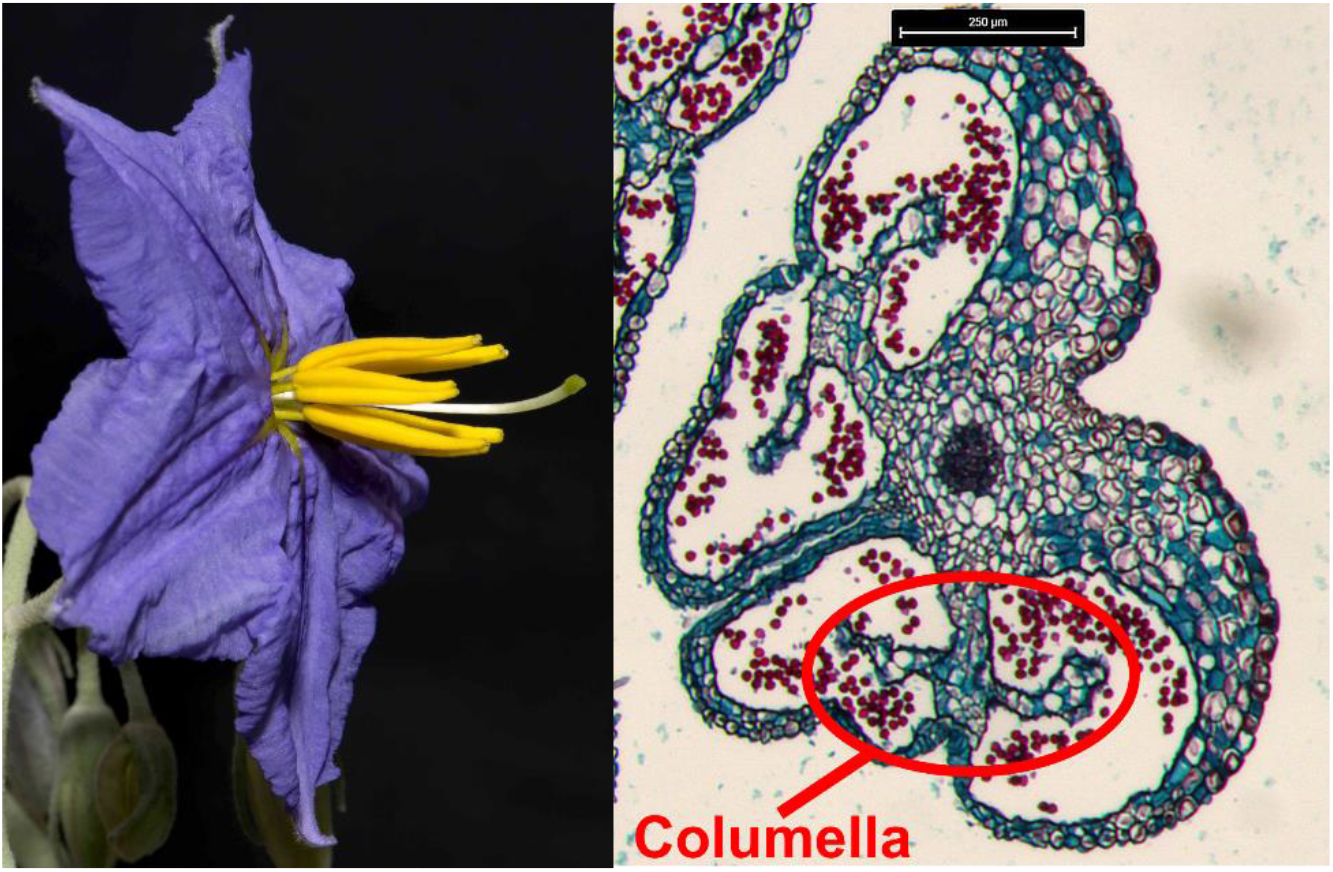
(Left) Image of a S. elaeagnifolium flower. The stamens (yellow) are composed of short cylindrical filaments supporting elongated poricidal anthers. (Right) Microtomed (20 μm thick) cross-section of a S. douglassii anther (which has nearly identical internal geometry when compared to S. elaeagnifolium), where the red dots indicate pollen grains located within the hollow anther locules. Note the internal columella circled in red on the anther cross section.

In addition to vibration frequency and amplitude, the spatial variation of vibration within the anther may also influence pollen expulsion. The pattern of deformation experienced by the vibrating anther affects how kinetic energy is imparted to the pollen grains and subsequently how the pollen grains exit the apical pore. Vibration transmission is thought to occur throughout different spatial axes within the flower and bees might maximize the stamen structural response by applying forces in directions associated with large structural amplification (see Nevard et al., 2021). For example, Nevard et al. (2021) showed that floral stamen bending may be more pronounced in one direction relative to another even when subject to a similar mechanical stimulus. However, this study considered only a single frequency excitation - but spatiotemporal vibration is highly dependent on the specific excitation frequency (Meirovitch, 2001). Intrinsically linked to each natural frequency of a continuous structure are its vibration mode shapes (Rao, 2019). Vibration mode shapes are the pattern of repetitive deformation a structure takes on when excited identically at the corresponding natural frequency. Deforming structures can often be described as a linear combination of their weighted vibration modes. The mode shapes, and how they contribute to the overall motion of the deforming stamen during a bees’ buzz, are therefore of fundamental importance to pollen excitation and expulsion from a poricidal anther. To date, they have not been characterized experimentally (but see M. J. King & L. Lengoc, 1993).

Beside physical experimentation, computational models could also be used to estimate the stamen’s vibration modes, natural frequencies, and physical deformations under specified loading conditions. Finite element analysis (FEA) is perhaps the most common approach to determining these quantities for structures with complex geometry (Reddy, 2005). FEA discretizes a continuous structure into smaller components (elements). External loads or prescribed motion are applied to the structure, and force equilibrium and displacement compatibility conditions are enforced at the interelement connections. Though approximate, FEA can estimate mode shapes and natural frequencies of the floral stamen where exact analytical representations cannot be formulated. In addition to estimating biophysical properties under normal conditions, FEA models could be used to explore stamen properties in situations that would be difficult or impossible to investigate experimentally. For example, an FEA model could show how material density and Young’s modulus affect stamen modes and natural frequencies, where Young’s modulus is defined as the material stiffness (Jastrzebski & Komanduri, 1988). Further, anthers have internal structures, such as the I-beam-like “columella” running along the axis of poricidal anthers (Figure 1; Halsted, 1890; van de Poel et al., 2014). FEA can be used to determine if or how such internal structures affect stamen dynamics. Lastly, an FEA model could quantitatively describe how the weight and location of a buzzing bee on the anther affects floral vibrations (e.g., Papaj et al., 2017; Solís-Montero & Vallejo-Marín, 2017). Despite these advantages, we know of only one other study that has used FEA to model a poricidal anther (M. J. King & L. Lengoc, 1993).

The goal of the present study is to experimentally characterize the natural frequencies, damping ratios and corresponding vibration mode shapes of a floral stamen. From morphometric and dynamic measurements, we develop a computational model of the stamen using FEA. We then use the FEA model to estimate how variable material properties, geometries and the mass of a bee located on the anther influence the stamen natural frequencies and modes. All studies are conducted on *Solanum elaeagnifolium,* a geographically widespread and common model organism in the study of floral buzzing that has elongated poricidal anthers and is visited by a variety of buzzing bees (Knapp et al., 2017, Figure 1).

## 2. Materials and Methods

### 2.1 Specimen Collection & Morphometrics

Seeds of *Solanum elaeagnifolium* Cav. collected 2.5 km NNE of Portal, Arizona (Cochise Co.) were scarified and grown in a Missouri State University greenhouse. Fresh flowers from 6-month-old plants were harvested on the day of anthesis and refrigerated until shipped later the same day. Flowers borne on short stems were placed in Aquatube florist water tubes within a plastic bag and shipped via FedEx to Bozeman, MT. Flowers were not older than 36 hours when received for experimentation. Individual stamens were removed from the flowers for experimental studies. The filament was left intact. Ablated stamens were weighed using a Mettler Toledo scale (XS603S, Mettler Toledo, Columbus, OH, USA) and were subsequently photographed on 5 mm x 5 mm gridded paper. Morphometric analyses were performed using ImageJ (vers. 1.53k). We measured the filament length, anther length, and maximum anther width.

### 2.2 Experimental Modal Analysis

We performed experimental modal analysis (EMA) to characterize the first several natural frequencies, damping ratios and mode shapes of the *S. elaeagnifolium* stamen (Figure 2). EMA for each specimen was conducted within fifteen minutes of removal from the flower to minimize the influence of desiccation on structural properties. The stamen was clamped in an inverted position by the proximal end of the filament (leaving the filament and anther free to move). Both sides of the clamp were lined with 3.2 mm hobby foam so that when closed, the hobby foam fully surrounds the filament base. This prevents circumferential crushing, which could potentially reduce the stamen’s natural frequencies. The clamp was attached to an electrodynamic vibration shaker with an amplifier rated at 31 N of force (K2007E007, Modal Shop, Sharonville, OH, USA). The shaker was excited via an external function generator with a linear chirp signal sweeping from 0 - 5000 Hz over a span of 640 milliseconds. During the linear chirp signal, the frequency increases continuously and linearly over the signal duration.

**Figure 2.**
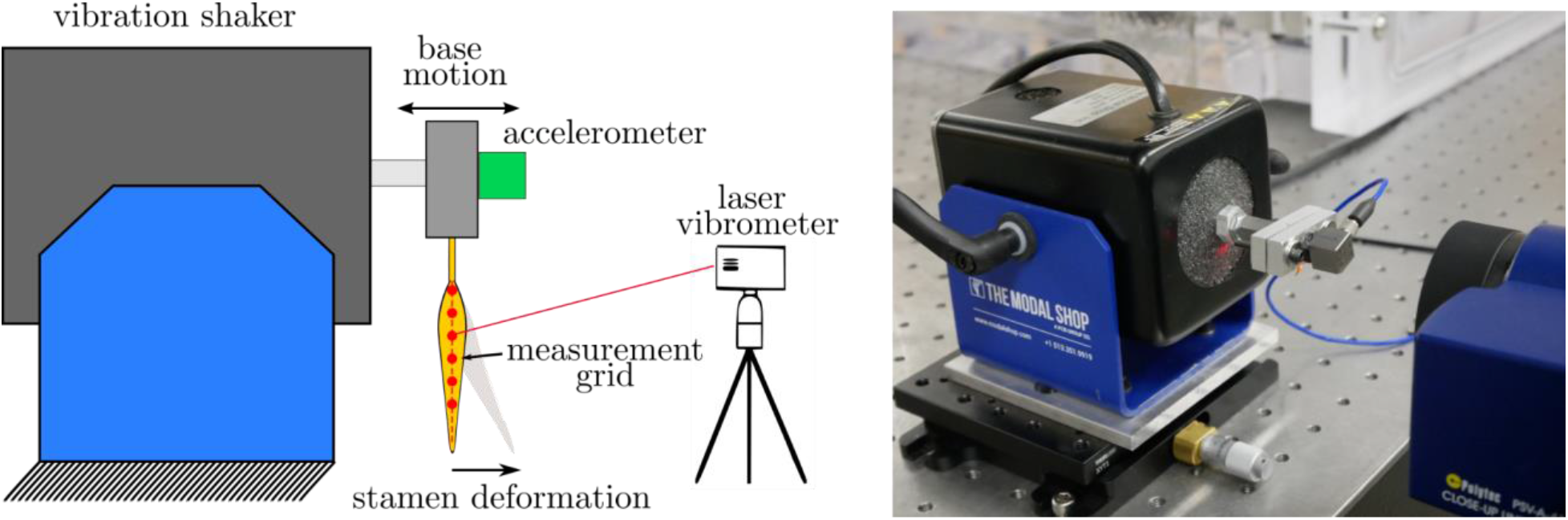
(Left) Schematic of experimental set-up, (Right) Physical realization. Schematic components are not drawn to scale.

The base motion of the shaker was measured by a high-sensitivity (500 mV/g) modal accelerometer (333B40, PCB Piezotronics, Depew, NY, USA). Base acceleration (considered the dynamic input to the stamen) was restricted to small accelerations between 2.6 mm-s^-2^ and 7.5 mm-s^-2^ across the frequency range considered to keep the stamen in the linear-elastic regime. This prevents permanent deformation of the stamen, which may alter its natural frequencies. The stamen output velocity was measured using a scanning laser vibrometer (PSV-400, Polytec, Hudson, MA, USA) with close-up attachment (PSV-A-410, Polytec, Hudson, MA, USA). Velocity was measured at 15 - 20 evenly spaced points along the midline of the anther from base to tip. The response at each measurement point was recorded three times and averaged. Base acceleration and stamen velocity data were acquired at a rate of 12.8 kHz, resulting in a frequency resolution of 1.5625 Hz across 3200 spectral lines over the frequency range considered.

Measured responses were reconstructed into operating deflection shapes using Polytec Presentation Viewer (Polytec, Hudson, MA, USA, vers. 9.3.1.3). When evaluated at the stamen natural frequencies, the operating deflection shapes are considered the vibration mode shapes. We used the local frequency response curve fitting algorithm to identify the stamen peak frequencies and damping ratios. In the case of light damping, the peak frequencies coincide closely with the stamen natural frequencies. The peak frequencies and mode shapes therefore serve as validation metrics for the stamen FE model developed in this work.

### 2.3 Finite Element Modeling

We developed a finite element model of the *S. elaeagnifolium* stamen from measured morphometric properties. A 3D model of the stamen was created using 35mm DSLR macro photographs of several stamens from different angles and magnifications as reference materials. Internal cross-sectional anatomy of the hollow anther locules was based upon photomicrographs of microtomed anther thin sections (20 μm) stained with Safranin and Fast Green from an earlier study (Buchmann et al., 1978). Thin sections allowed us to see the extent of the connective tissue (columella) that extends into each of the paired hollow anther locules as a ridge extending along their full length. The 3D stamen was modeled by a professional 3D Tucson artist using ZBrush vers. 2020. The artist also developed a half-length version, and a version without a columella so that we could study how these geometric properties affected natural frequencies (Figure 3).

**Figure 3.**
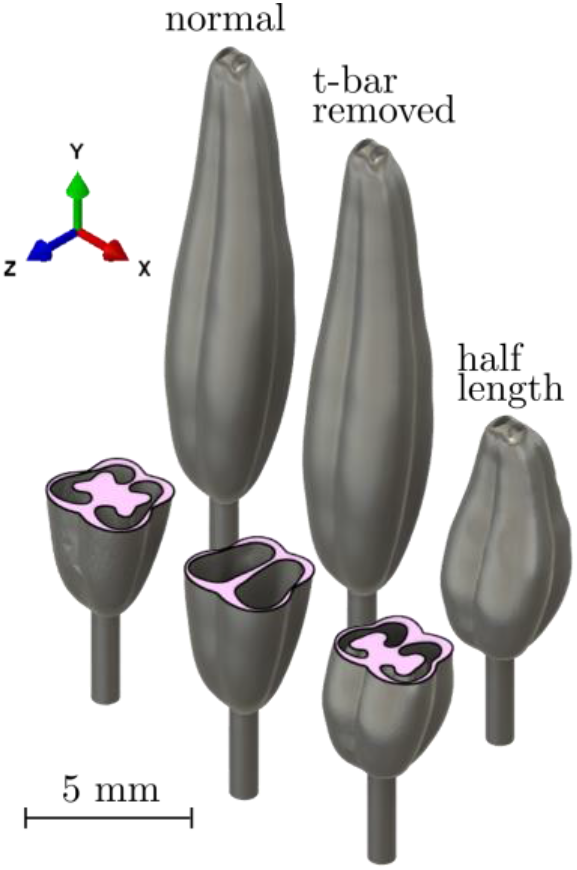
3D renderings of the geometric .stl models used for our FE studies. The furthest left image shows the normal anther, whereas the right two images show the anther’s geometric modifications. The center anther has the internal columella removed, and the right anther is compressed so that it has half the length of the normal anther.

The 3D stamen geometry was imported into ABAQUS (Dassault Systemès, Vélizy-Villacoublay, France, Ver. 6.23) and scaled to match the 16.7 mm average stamen length (sum of average filament and anther lengths, Table 1). We assumed the stamen to have linear-elastic and isotropic material properties. Material density was set to 154 kg-m^-3^ so that the stamen model had equivalent mass to the average measurement of 5.96 mg. This density value is similar to the bulk density values reported for *Crocus sativus* L. anthers (Emadi & Saiedirad, 2011). Young’s modulus was set to 11.88 MPa such that the first calculated natural frequency coincided with the first measured natural frequency. While Young’s modulus has not been reported for *S. elaeagnifolium* stamen tissue, the 11.88 MPa value falls within the reported range for other plant tissues (Gibson, 2012). Poisson’s ratio (the ratio between a material’s transverse and axial strains) was approximated as 0.45 (only known for pumpkin tissue; Shirmohammadi et al., 2018). The model was meshed into 213,127 quadratic tetrahedral elements (C3D10), which was sufficient for convergence of the first six calculated natural frequencies. We fixed all degrees of freedom at the base of the filament. The first six natural frequencies and corresponding mode shapes were determined using ABAQUS. Quantities calculated via FEA were compared to our experimental measurements to validate the accuracy of the FEA model.

**Table 1.** Summary of morphometric parameters of stamen (n = 19)

|  | **Filament Length (mm)** | **Anther Length (mm)** | **Max Width (mm)** | **Mass (mg)** |
| --- | --- | --- | --- | --- |
| **Mean** | 3.05 | 13.65 | 2.29 | 5.96 |
| **Std. Deviation** | 0.78 | 1.46 | 0.27 | 1.33 |
| **First Quartile** | 2.67 | 12.70 | 2.16 | 5.63 |
| **Third Quartile** | 3.55 | 14.87 | 2.44 | 6.67 |

Once the FEA model was validated, we performed a series of parametric studies to characterize how variable stamen tissue properties, morphologies and external loading conditions affected the stamen natural frequencies and vibration modes. First, we characterized the natural frequencies as a function of Young’s modulus and density. We considered five evenly spaced Young’s modulus values in the range 6 - 18 MPa and five evenly spaced density values in the range 77-231 kg-m^-3^, where the lower and upper bound of the range are determined as 50% and 150% of the normal parameter value. Next, we investigated the modal characteristics of two hypothetical stamen: one stamen where the anther was 50% reduced in length, and a second full-length anther where the columella was removed. The same 3D mesh density as the full-length anther was maintained for these alternative geometries. Finally, we studied the effect that bee mass has on stamen dynamic properties. We considered both the location of the bee on the anther and the mass of the bee. First, we treated the bee as a point mass of 150 milligrams located 0%, 25%, 50% and 75% from the tip of the anther in separate numerical studies. Next, we fixed the bee location at 50% from the anther tip and varied the bee mass from 50 – 450 mg in 100 mg increments, where the range was selected to encompass multiple species of floral buzzing bees (Bullock, 1999; Cane, 1987; Hagen & Dupont, 2013; Jankauski et al., 2021). Gravity (g = 9.81 m-s^-2^) acts in the positive z direction (Figure 3). We first solved for the static deformed shape of the stamen using the Newton-Raphson method which accounts for the geometric nonlinearity associated with large bending at the filament. The Newton-Rhapson solver within ABAQUS is its default nonlinear structural solver and is often used to solve nonlinear static structural problems (Stricklin & Haisler, 1977). We subsequently performed numerical modal analysis about the statically deformed shape. To facilitate calculation during the nonlinear static step, we reduced the anther mesh to 16,562 from 213,127 elements for this parametric study only. Ignoring the point mass and gravity, the difference between the six stamen natural frequencies calculated using the original fine mesh and the modified coarse mesh was only 0.78%.

## 3. Results

### 3.1 Morphometrics

We collected morphometric data from n=19 stamens collected from five *S. elaeagnifolium* flowers. Stamen morphometric parameters are summarized in Table 1. The stamen had a mean filament length of 3.05 mm, a mean anther length of 13.65 mm, a mean maximum width of 2.29 mm, and a mean mass of 5.96 mm.

### 3.2 Experimental Modal Analysis

We used experimental modal analysis (EMA) to identify *S. elaeagnifolium* stamen (n=7 stamens from two flowers) natural frequencies, damping ratios and vibration mode shapes. A representative frequency response magnitude plot from which natural frequencies and damping ratios are extracted is shown in Figure 4, and determined quantities are summarized in Table 2. Over the experimental excitation range, we identified two natural frequencies and vibration modes. The first mode occurred at about 64.2 Hz and corresponded to an out-of-plane bending mode (Figure 5), where out-of-plane modes are characterized by motion predominantly in the z-direction. The second mode occurred at 1227.9 Hz and corresponded to an out-of-plane mode where the base and the tip of the stamen move antiphase with respect to one another and are separated by a stationary nodal point (Figure 5). The first and second modes had damping ratios of approximately 0.12 and 0.085 respectively, which indicates that the stamen are relatively underdamped. These average natural frequencies and mode shapes serve as validation metrics for the FEA model described in the following section.

**Figure 4.**
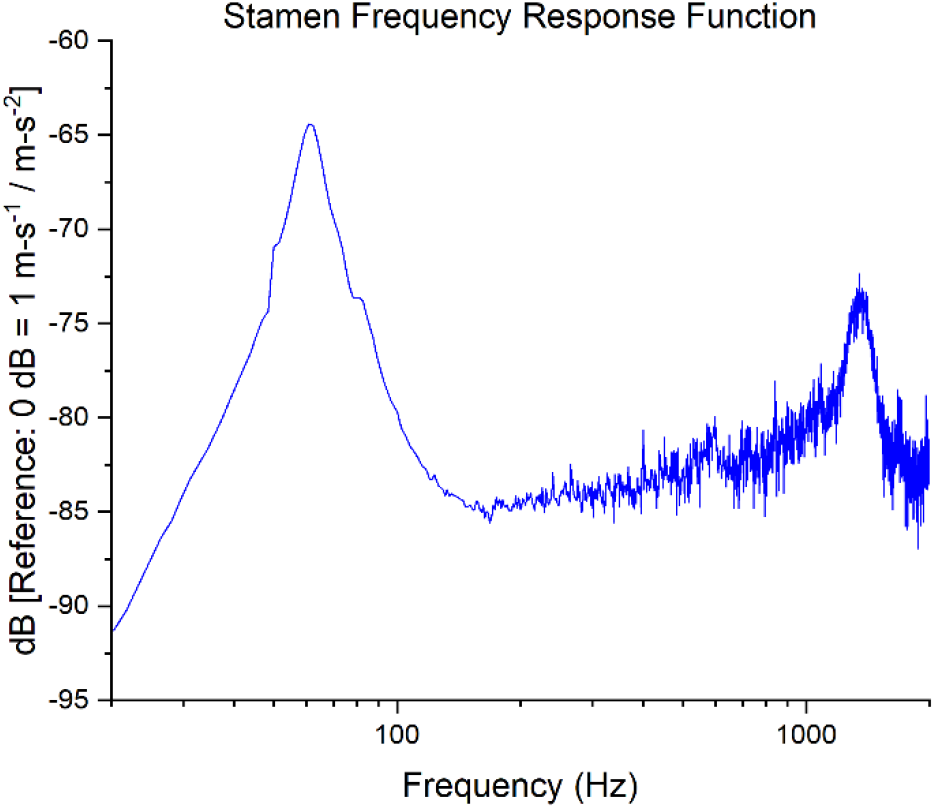
Representative frequency response function from which stamen natural frequencies and damping ratios are determined. Note the resonant peaks around 60 and 1200 Hz.

**Table 2.**
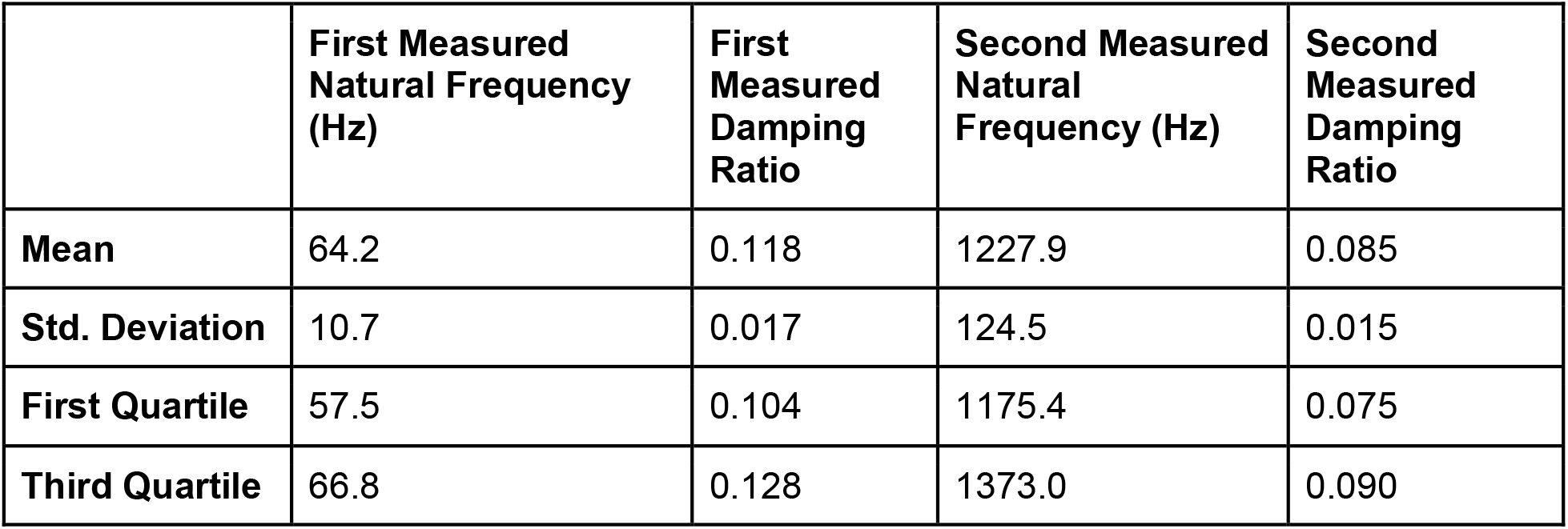
Summary of stamen dynamic properties (n = 7)

|  | **First Measured Natural Frequency (Hz)** | **First Measured Damping Ratio** | **Second Measured Natural Frequency (Hz)** | **Second Measured Damping Ratio** |
| --- | --- | --- | --- | --- |
| **Mean** | 64.2 | 0.118 | 1227.9 | 0.085 |
| **Std. Deviation** | 10.7 | 0.017 | 124.5 | 0.015 |
| **First Quartile** | 57.5 | 0.104 | 1175.4 | 0.075 |
| **Third Quartile** | 66.8 | 0.128 | 1373.0 | 0.090 |

**Figure 5.**
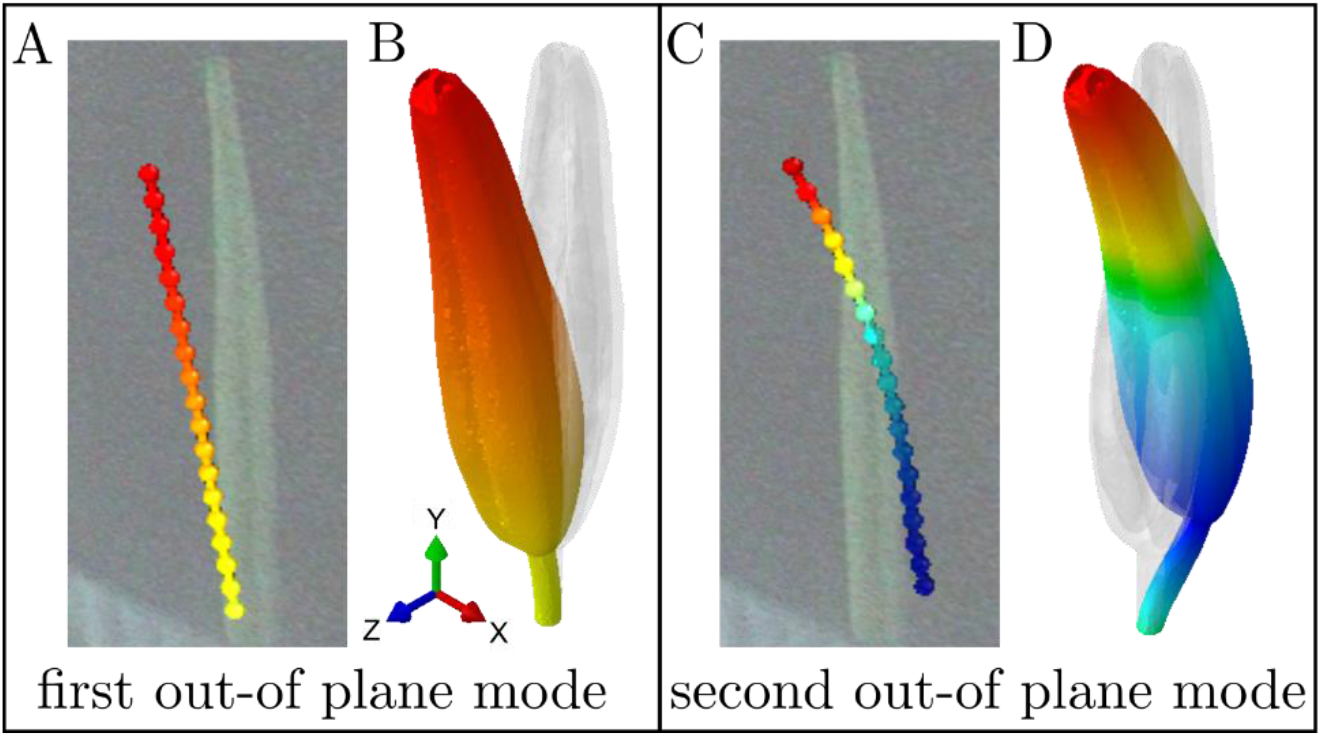
Comparison between measured (A,C) and calculated (B,D) vibration modes. Individual dots in panels A,C indicate the vibrometer measurement points along the anther. Yellow indicates regions of no motion, red indicates denote regions with motion in the +z direction, and red indicates regions with motion in the -z direction.

Our experimental setup may not capture all natural frequencies and vibration modes within the experimental excitation range. The laser vibrometer is capable only of measuring out-of-plane motion and consequently in-plane modes may not be observed. In-plane modes are characterized by motion in the x-direction (Figure 3). Given the near symmetry of the stamen (particularly at the filament), modes dominated by bending likely appear in pairs with identical or closely spaced natural frequencies (see example in Wake et al., 1998). We hypothesize that there are unmeasured in-plane modes similar to each of the out-of-plane modes shown in Figure 5. Note that even if vibration modes and natural frequencies occur in pairs, they must still be considered distinct. Furthermore, due to stamen symmetry and prescribed base motion during EMA, we may not excite torsional modes where the stamen twists about its longitudinal axis. Torsional modes would only be excited if the stamen center of mass did not coincide with its longitudinal axis or a direct force was applied to the stamen away from the longitudinal axis.

### 3.3 Finite Element Model Validation

Here, we compare the natural frequencies and mode shapes measured experimentally to those predicted computationally. The first and second measured natural frequencies were 64.2 Hz and 1227.9 Hz respectively, which are close to the natural frequencies of 64.2 Hz and 1234.4 Hz predicted by FEA for similar vibration modes. The Young’s modulus used in the model was adjusted so that the first calculated natural frequency aligned with experimental results, so the difference between the second reported natural frequency better represents model error.

Measured and estimated vibration mode shapes corresponding to these natural frequencies are shown in Figure 5. There are no notable discrepancies between measured and predicted modes. The first mode corresponds to the first out-of-plane bending mode where the stamen rocks back and forth and experiences maximum deflection at its tip. The second mode corresponds to the second out-of-plane bending mode, where the base of the stamen moves antiphase to its tip. With confidence that the FEA model captured the real stamen natural frequencies and mode shapes, we then used the model to better understand how different parameters influenced stamen dynamics through a series of parametric studies.

### 3.4 Finite Element Parametric Studies

The stamen natural frequencies and mode shapes calculated via FEA are shown in Figure 6. The computational results indicate several modes that were not captured experimentally. Each bending mode occurred in out-of-plane/in-plane pairs with nearly identical natural frequencies. The calculated natural frequencies differ only due to the asymmetry of the anther; if the anther were cyclically symmetric, we would expect these natural frequencies to coincide exactly. Between the two pairs of bending modes, we observed a torsional mode where the stamen twists along its longitudinal axis. The sixth mode corresponded to an axial mode, where the stamen stretches and compresses along its longitudinal axis like a coil spring.

**Figure 6.**
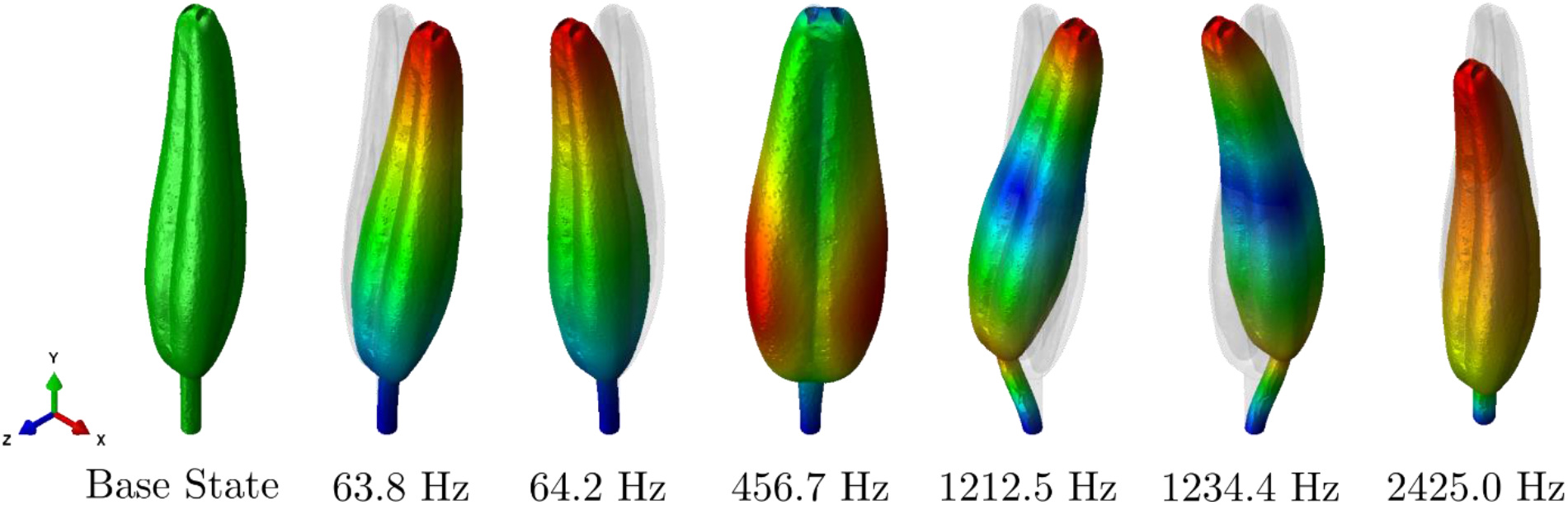
Undeformed state of the anther (far left) and first six vibration modes from left to right. Blue indicates zero modal displacement and red indicates maximum modal displacement. Color scale bar not included, since vibration mode shapes are arbitrary in magnitude and modal displacements arbitrary in units.

Variation of Young’s modulus and density influenced the stamen natural frequencies but not its mode shapes. These properties influenced all natural frequencies proportionally, meaning if a deviation of Young’s modulus and density doubled the natural frequency of one mode, it would double the natural frequencies of all other modes as well. The ratio of the modified natural frequency to the original natural frequency 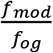 is shown as a function of Young’s modulus and density in Figure 7. Similar to a fixed-free cantilever beam, the stamen natural frequencies scale linearly with the square root of the Young’s modulus 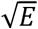 and with one over the square root of density 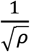 (Rao, 2019). Therefore, we can use a simple bilinear regression model to identify 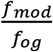 as

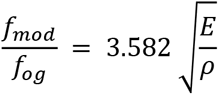

where *E* is the Young’s modulus in MPa and *ρ* is the density in kg-m^-3^.

**Figure 7.**
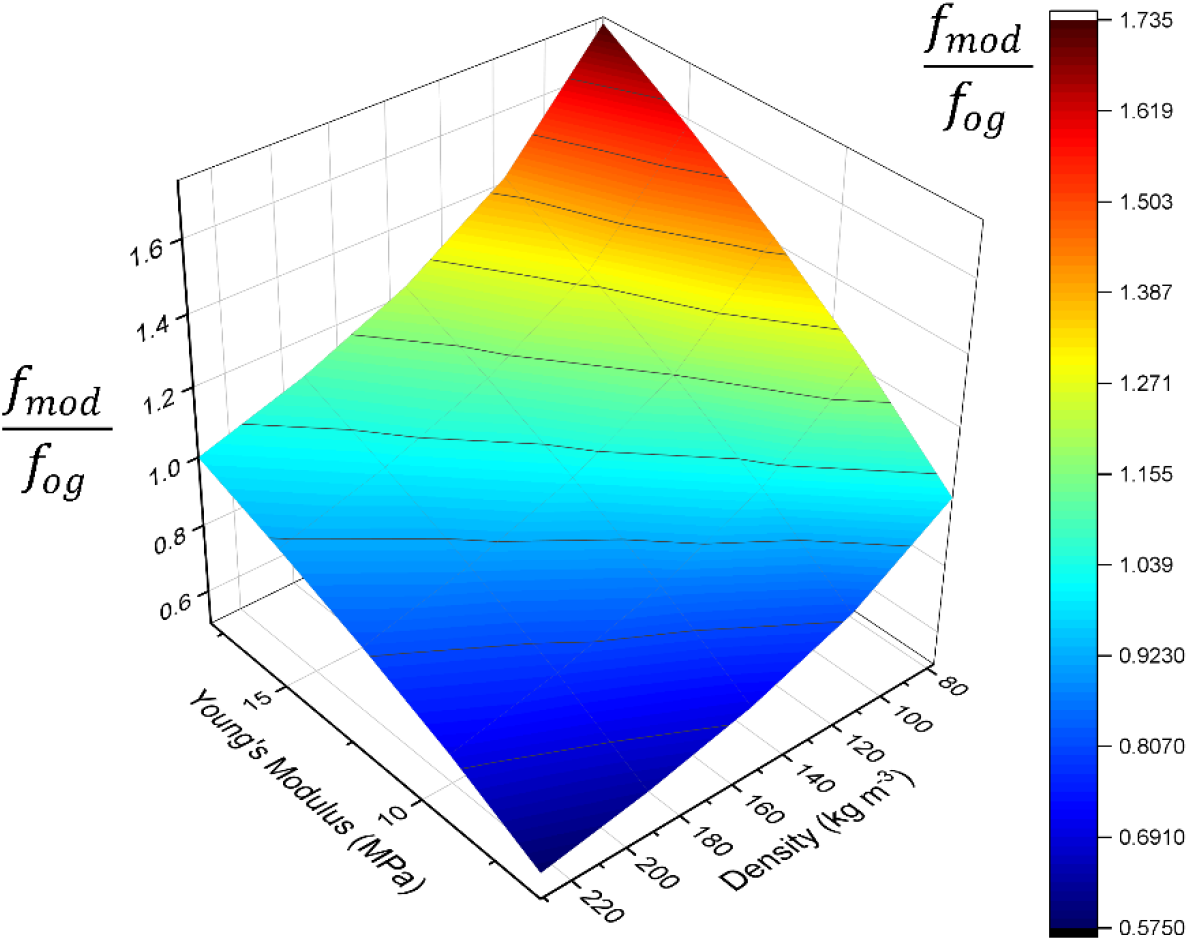
Surface plot showing the influence of Young’s modulus and density on the stamen natural frequency. f denotes the original natural frequency calculated using a Young’s modulus of 12 MPa and a density of 154 kg^-3^. f_mod_ denotes the modified natural frequency calculated using an alternative set of Young’s modulus and density values as specified by the grid.

Altering the stamen geometry influenced its natural frequency but did not significantly affect the characteristics of the vibration mode shapes. The first six natural frequencies of the normal stamen, stamen with columella removed, and half-length stamen are shown in Table 3. Interestingly, removal of the columella caused all natural frequencies to increase slightly. In general, the columella may increase the stiffness of the anther, but also increases the anther mass, which in turn reduces the natural frequency of the entire stamen. The columella would increase the natural frequency only if it extended through the filament and was fixed at the base. Nonetheless, the columella structure likely increases the bending rigidity of the anther itself, perhaps so that the anther pores do not collapse and preclude pollen expulsion during floral buzzing. Reducing the anther length by 50% increased all natural frequencies. We expect this result since cantilever beam bending modes scale with 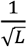, where *L* is the length of the beam (Rao, 2019). The reduced length anther does not quadruple the bending natural frequencies since the filament length is maintained between models.

**Table 3.** Summary of stamen natural frequencies for geometric modifications.

|  | **$f_1$ (Hz)** | **$f_2$ (Hz)** | **$f_3$ (Hz)** | **$f_4$ (Hz)** | **$f_5$ (Hz)** | **$f_6$ (Hz)** |
| --- | --- | --- | --- | --- | --- | --- |
| **Normal** | 63.8 | 64.2 | 456.7 | 1212.5 | 1234.4 | 2425.0 |
| **Columella Removed** | 74.4 | 77.0 | 517.3 | 1385.3 | 1406.5 | 2826.7 |
| **Half Length** | 151.7 | 152.0 | 647.3 | 1926.4 | 1991.7 | 3560.7 |

We now direct our attention to the influence the bee has on the stamen. The weight of the bee causes the stamen to bend substantially. The maximum anther tip deflection is 1.78 mm when the 150 mg bee is located 75% from the tip of the anther and 5.10 mm when the mass is located 0% from the tip of the anther (Figure 8). When the bee location is fixed at 50% down the length of the anther and mass is varied, the anther tip displaces 1.03 mm for a 50 mg bee mass and 7.25 mm for a 450 mg bee mass. Unlike the other parameter studies, incorporating the weight of the bee into the FEA model influenced both stamen natural frequencies (Tables 4,5) and most of the vibration mode shapes. For all cases, the first two modes corresponded to in-plane and out-of-plane bending modes but occurred at lower natural frequencies compared to the unweighted stamen. Higher-order modes diverged from those observed in the unweighted stamen but had similar characteristics.

**Figure 8.**
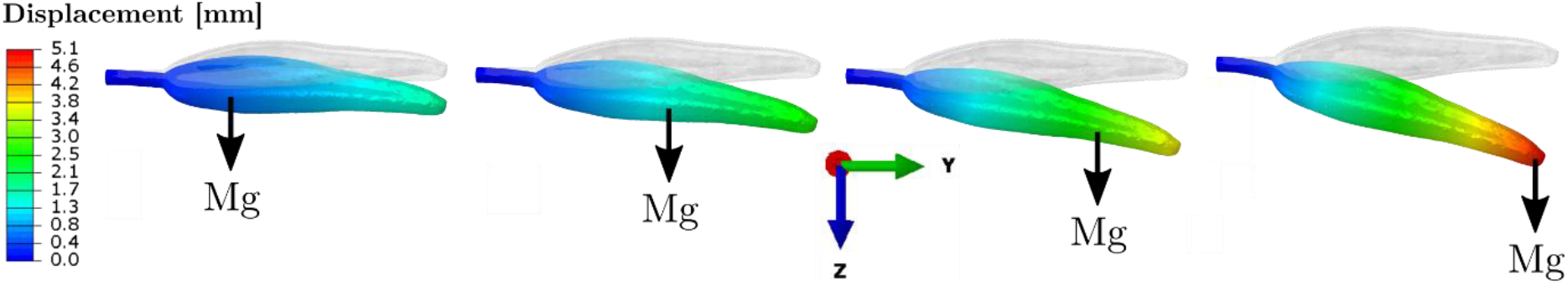
Static deformation calculated via FEA for a stamen with gravity and bee mass (M = 150 milligrams). Simulations are conducted assuming the bee is attached (from left to right) 25%, 50%, 75% and 100% from the base of the anther. ‘Mg’ denotes the force due to gravity, where ‘M’ is the anther mass and ‘g’ is gravitational acceleration.

**Table 4.** Summary of stamen natural frequencies when bee mass and gravity are considered, and bee mass is fixed at 150 mg. Location of the bee is varied as a percentage of anther length between 0 – 75% from the anther tip. The brown highlighted cells show the natural frequencies corresponding to the mode we hypothesize is excited (Figure 9) by the bee during floral buzzing.

| Point Mass Location (%) | $f_1$ (Hz) | $f_2$ (Hz) | $f_3$ (Hz) | $f_4$ (Hz) | $f_5$ (Hz) | $f_6$ (Hz) |
| --- | --- | --- | --- | --- | --- | --- |
| 0 | 7.0 | 7.2 | 239.1 | 420.2 | 826.9 | 863.7 |
| 25 | 9.1 | 9.2 | 287.7 | 391.7 | 1115.6 | 1206.2 |
| 50 | 12.8 | 13.0 | 312.6 | 314.0 | 1181.6 | 1284.7 |
| 75 | 19.4 | 19.7 | 201.1 | 268.9 | 725.1 | 978.4 |

**Table 5.** Summary of stamen natural frequencies when bee mass and gravity are considered, and bee location is fixed at 50% of the anther length from the anther tip. Bee mass is varied between 50 – 450 mg. The brown highlighted cells show the natural frequencies corresponding to the mode we hypothesize is excited (Figure 9) by the bee during floral buzzing.

| Bee Mass (mg) | $f_1$ (Hz) | $f_2$ (Hz) | $f_3$ (Hz) | $f_4$ (Hz) | $f_5$ (Hz) | $f_6$ (Hz) |
| --- | --- | --- | --- | --- | --- | --- |
| 50 | 21.1 | 21.2 | 318.0 | 562.1 | 1184.9 | 1238.1 |
| 150 | 12.8 | 13.0 | 312.6 | 314.0 | 1181.6 | 1284.7 |
| 250 | 10.2 | 10.4 | 230.0 | 311.9 | 1179.0 | 1343.9 |
| 350 | 8.9 | 9.1 | 183.9 | 311.7 | 1177.0 | 1395.7 |
| 450 | 8.1 | 8.4 | 154.1 | 311.4 | 1175.1 | 1437.7 |

Notably, the bee weight lowers the natural frequency of the axial vibration mode and introduces an out-of-plane motion to this mode as well. This new axial-bending vibration mode (Figure 9) occurs regardless of bee mass value and location. The brown cells highlighted in Tables 4 and 5 show the natural frequencies corresponding to this mode for each bee mass value and location. The natural frequencies for the axial-bending mode range from 154.1 - 562.1 Hz, which coincides with the buzz frequency of many floral buzzing species (De Luca & Vallejo-Marín, 2013; Arroyo-Correa et al. 2019; Switzer et al. 2019; and references within). Thus, we believe this mode may contribute considerably to the stamen motion during floral buzzing.

**Figure 9.**
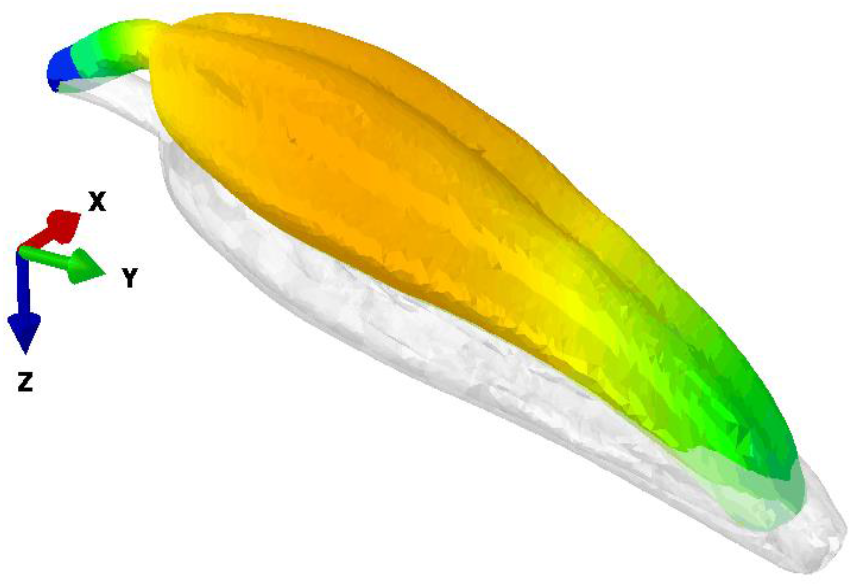
New axial-bending mode that arises when the bee mass and gravity is considered. The natural frequencies corresponding to this mode occur between 154.1 - 562.1 Hz depending on where the bee mass magnitude and location. This frequency range coincides with reported floral buzzing frequencies. Blue indicates zero modal displacement and red indicates maximum modal displacement. Color scale bar not included, since vibration mode shapes are arbitrary in magnitude and modal displacements arbitrary in units.

## 4. Discussion

### 4.1 Natural Frequencies

Exciting a structure near one or more of its natural frequencies generally elicits a large vibrational response due to resonance (De Langre, 2019), which in the case of poricidal anthers, could result in enhanced pollen expulsion. Thus, a long-standing question in the study of poricidal flowers buzzed by bees is why the natural frequency of stamens falls outside of the buzzing frequency range for most bees, which has been reported from about 100 - 400 Hz (Arroyo-Correa et al., 2019; Burkart et al., 2012; De Luca et al., 2014, 2019; De Luca & Vallejo-Marín, 2013; Nunes-Silva et al., 2013; Switzer et al., 2016, 2019, see also Harder & Barclay, 1994). We found at least two *S. elaeagnifolium* stamen natural frequencies that occurred within the 0 - 5000 Hz experimental excitation range considered. The first occurred in the range between 49.8 – 85.9 Hz. This is lower than the reported range of approximately 100 – 145 Hz for *Solanum laciniatum* (King & Buchmann, 1996) but within the 45 – 295 Hz range reported for other *Solanum* taxa (Nunes et al., 2021). The second measured natural frequency for *S. elaeagnifolium* stamens was in the range 1175 - 1375 Hz. As with prior studies, these stamen natural frequencies and those predicted by finite element analysis occurred outside the floral buzzing frequency range for most bees. While variable geometry or material properties affect the stamen natural frequencies to some degree, these variations themselves are unlikely to raise the first measured natural frequency or lower the second measured natural frequency to within the range of bee floral buzzes.

By contrast, the weight of a medium to large bee (such as a bumblebee or carpenter bee) dramatically shifts the stamen natural frequencies. Deviations in natural frequencies depend on the weight of the bee as well as where the bee’s center of mass is located on the stamen. Floral buzzing bees may weigh 20 - 100 times more than the weight of the stamen they are visiting to collect pollen (see Bullock, 1999; Cane, 1987; Hagen & Dupont, 2013; Jankauski et al., 2021). Natural frequencies are inversely proportional to mass, and so this considerable shift in mass dramatically influences stamen dynamics. Masses located further from the fixed stamen base influence the natural frequencies more than masses located near the fixed stamen base. Our FEA simulation suggests that the third and fourth stamen natural frequencies occur between 154.1 – 562.1 Hz depending on the location and mass of the bee. This range is substantially more consistent with the reported floral buzzing frequency range of bees. Thus, we hypothesize that floral buzzing bees do exploit stamen resonance or near resonance to generate a large structural response of the anther. This may facilitate dislodging of pollen, pollen-to-wall interactions, and the subsequent expulsion of pollen through the two small apical anthers pores in *S. elaeagnifolium.* This finding also suggests a mechanism governing bee specialization among poricidal plant species; to maximize pollen collection, bees should specialize on flowers that match stamen resonant properties, given the bee’s weight (e.g., Corbet & Huang, 2014). However, previous studies have indicated that pollen expulsion may be more correlated to vibration amplitude than frequency in some poricidal anthers (De Luca et al., 2013; Rosi-Denadai et al., 2020). It may therefore be possible to expel pollen with forces that are sufficiently high in magnitude but at a frequency mismatched from the stamen’s resonant frequency.

### 4.2 Vibration Mode Shapes

Mode shapes describe the vibratory pattern a structure follows if excited at one of its natural frequencies and are of fundamental importance to pollen excitation and expulsion from a poricidal anther. The vibration mode shapes of a stamen are like those of a cantilever beam. Numerical simulation shows the stamen has in-plane and out-of-plane bending modes, as well as torsional and axial modes. Excitation of a mode depends both on the proximity of the input forcing frequency to the mode’s natural frequency, as well as the location at which the force is applied. To excite a vibration mode, the periodic force should be applied at a location where the modal displacement is greatest.

Interestingly, the axial-bending mode (Figure 9) present when the bee weight is considered satisfies both criteria. This mode has natural frequencies between 154.1 – 562.1 Hz (dependent on bee mass magnitude and location), which falls within the reported buzz frequency range for many species of bees (Arroyo-Correa et al., 2019; Burkart et al., 2012; De Luca et al., 2014, 2019; De Luca & Vallejo-Marín, 2013; Nunes-Silva et al., 2013; Switzer et al., 2016, 2019). Further, its modal displacement is greatest in the transverse direction about 60% down from the anther tip. This is close to where floral buzzing bees typically bite the anther (Macior, 1964; Papaj et al., 2017; Switzer et al., 2016), and where the periodic load resulting from flight muscle vibration is effectively applied. Recent studies suggest the largest forces generated by defensively buzzing carpenter bees act in a direction transverse to the anther (Jankauski et al., 2021). Though there is some debate to whether defensive buzzing is biomechanically similar to floral buzzing (see De Luca et al., 2014; Pritchard & Vallejo-Marín, 2020), we expect forces would be applied in the same direction during both behaviors. We thus hypothesize that if the axial-bending mode is excited, the large lateral motion of the locule walls would impart significant amounts of kinetic energy to the enclosed pollen grains, and the axial motion of the locule would produce a frictional force, moving the pollen towards the apical pore.

### 4.3 Computational Modeling

To date, there have been few efforts to develop a high-fidelity explanation or model of a poricidal stamen capable of describing its structural dynamics. While some models describe pollen ejection, the anther cavity and cavity wall motion in these models is greatly simplified (Buchmann & Hurley, 1978; Hansen et al., 2021). The present work is particularly useful when simulating conditions difficult to emulate and study experimentally under controlled laboratory conditions. For example, during floral buzzing, the bee may contact a large portion of the anther with its body or its legs may contact other parts of the flower. This makes it challenging to measure the physical displacement of the anther using laser vibrometry or high-speed videography. Further, a computational model allows one to pose potential scenarios to identify how the dynamics of the stamen may change under different circumstances, such as variable material properties and how different geometric features contribute to stamen dynamics.

More importantly, this structural model can be integrated with a pollen dynamics model to better understand how pollen is expelled from the vibrating anther. However, additional studies must be conducted to understand the coupling factors between the hollow anther locules and the thousands of pollen grains within. If pollen mass is small compared to anther mass, it may be possible to simulate the structural response of the anther while ignoring the pollen mass. However, at 25% of the anther mass (such as for *S. elaeagnifolium,* with about 1 mg of pollen per anther; Russell & Buchmann, unpublished data), pollen mass may influence stamen natural frequency. Indeed, experimental studies conducted on wind pollinated *Plantago lanceolata* have shown that stamen natural frequency increases as pollen is shed from the vibrating anther (Timerman et al., 2014). If this is also the case for poricidal flowers, the structural and pollen models would have to be solved in parallel, since the moving locule boundary would depend on the amount of remaining pollen. Shifts in natural frequency due to pollen mass loss may also underlie a pollen dosing strategy, since the stamen vibration amplification factor would diminish if the bee maintained a constant floral buzzing frequency. Finally, and critically, our work suggests that studies investigating the influence of floral buzz frequencies on pollen expulsion must consider the key influence of bee weight on stamen resonant properties.

## Acknowledgments

Thanks to Daniel Hornung for assistance creating the 3D stereo lithography models of the *S. elaeagnifolium* anthers. Buchmann thanks Sammy Missoum (Univ. AZ Dept. of Aerospace and Mechanical Engineering) for discussions about vibration along with an earlier FEA Solanum model. We thank C.E. Jones and D. R. Papaj for their comments about *S. elaeagnifolium* and buzz pollination over many years. Buchmann thanks Jack H. Burk for the use of his CSUF microtome and histology laboratory.

